# Arthropod Diversity at Two Sites with Different Disturbance Levels at the Texas A&M International University

**DOI:** 10.1101/2024.10.01.616158

**Authors:** Eduardo Siller

## Abstract

This research aimed to measure arthropod species richness in two different sites with different disturbance levels in the Texas A&M International University campus to determine the overall biological diversity of the two sites and compare these measurements. As part of the research and methods, the Paleontological Statistics (PAST) software was used to calculate statistical indices in ecology that can help determine the difference between East and South sites. Some indices employed were the Shannon’s Diversity Index or the Simpson’s Index of Diversity, the Bray-Curtis matrix, and the iNaturalist app. Results showed that the abundance of arthropods differed significantly from one site to the other, with the South site having a greater quantity, which was the site with less degradation created by humans. The site with the degraded ecosystem (East site) did not show a clear disadvantage regarding the diversity of arthropods. Thus, this article is relevant considering abundance and diversity to discover which is more important in each location. Finally, the study provides more insight into the arthropod community in the South Texas region, which can help with land and ecosystem management.

## Introduction

The phylum Arthropoda is one of the most diverse in the animal world. In it, one can find very different-looking organisms, from spiders, centipedes, and scorpions to bees, shrimps, and many more. Arthropods constitute more than 85% of all animal species, with over 5 million different species occupying almost every ecosystem of every continent (Goddard, 2018).

Because of their abundance and diversity, they are a vital part of any ecological community with many crucial roles. The leading roles of arthropods include pollinators, decomposers, and the service they provide as biological control of pests. Arthropods are essential in every ecosystem, and arthropod diversity and species richness must be adequately managed. Furthermore, arthropod diversity and species richness can measure an ecosystem’s overall diversity and quality (Obrist & Duelli, 2010).

The South Texas region is characterized by its semi-arid climate with little precipitation throughout the year, with an average of 62 cm of yearly rainfall (Atkinson *et al*., 2009). In addition, the region has hot, humid summers and mild winters, with average temperatures of 37°C and 5°C, respectively. That makes it an ideal ecosystem for certain types of desert-like flora and fauna, including several species of reptiles and cacti, such as the checkered garter snake (*Thamnophis marcianus*) and Texas prickly pear cactus (*Opuntia engelmannii*). Also, the South Texas region has many species of shrubs and trees, which compose an essential part of the vegetation of this area. These include native and nonnative species, such as honey mesquite (*Prosopis glandulosa*), black brush acacia (*Vachellia rigidula*), coyotillo (*Rhamnus humboldtiana*), Christmas cholla (*Cylindropuntia leptocaulis*), among others (Ruthven *et al*., 2002).

The Texas A&M International University (TAMIU) campus is located in Laredo, Texas, on the Southern border of the United States, along the Rio Grande River. The campus has a wide variety of species of plants and animals in the surroundings. The fauna around campus includes mammals, reptiles, amphibians, birds, arachnids, and insects, amongst others. These include species such as white-tailed deer (*Odocoileus virginianus*), Mediterranean house gecko (*Hemidactylus turcicus*), Texas toad (*Anaxyrus speciosus*), green jay (*Cyanocorax yncas*), striped bark scorpion (Centruriodes vittatus), and pipevine swallowtail (*Battus philenor*). By far, the most diverse are the arthropods, which include different species of butterflies, moths, scorpions, spiders, walking sticks, ants, crickets, ticks, among others.

This research aimed to measure arthropod species richness in two different sites with different disturbance levels in the TAMIU campus to determine the overall biological diversity of the two sites and make a warning regarding which site has been more affected and with propensity to endanger the species. Arthropods have been previously studied in different ecosystems due to their high diversity and abundance to determine the effect of habitat degradation on the organisms (Thibault *et al*., 2006). Thus, we followed an approach to determine the relation, or lack thereof, between arthropod diversity and habitat disturbance..

## Materials and methods

The study was conducted during June and July 2023 in the areas surrounding TAMIU (latitude 27.5714, longitude -99.4351). Due to the different degrees of human-caused ecosystem degradation, two different sites were selected in the east and south parts of campus for the study.

The first site, East of the TAMIU campus (latitude 27.5720, longitude -99.4285), is behind the dormitories. It is considered a degraded site as it has been the site of a tennis court complex construction in the last few years. The ecosystem has suffered habitat destruction; however, there is enough vegetation in the surrounding areas to collect data on arthropod diversity and abundance. The second site, on the South part of campus (latitude 27.5673, longitude -99.4330), is located across the soccer field. It was selected due to the low habitat loss and human disturbance.

The primary technique was the tree-shaking which used a beating stick and a beating sheet to collect arthropods from shrubs, bushes, and small tree branches (Muelelwa *et al*., 2010). This method was performed for 30 seconds to maximize the amount of arthropods. A white beating sheet is placed under the tree branches to gather arthropods. After 30 seconds, the arthropods that landed on the sheet are counted, recorded, and identified using the mobile app iNaturalist (version 2.8,6). The iNaturalist app is an artificial intelligence database that gathers information from hundreds of thousands of observations to suggest a species with 90% accuracy that can be verified by scientists worldwide based on the photographs taken. Data was collected weekly, randomly selecting five trees or shrubs for sampling from each site. The sampling occurred between 3:00 PM and 7:00 PM on the same day every week.

Once all data was collected, it was analyzed using Shannon’s Diversity Index, Simpson’s Index of Diversity, and Simpson’s Reciprocal Index to evaluate each site’s species richness, abundance, and evenness. Shannon’s index is a mathematical measurement that anticipates a randomly chosen species from a community (Cox, 2002). A proportion is calculated based on the number of individuals of each species and the total number of individuals of all species.

Simpson’s Index considers the number of species and the abundance of each species to represent the probability that two randomly selected individuals from a community belong to all species (Cox, 2002). The values range from 0 to 1, where 0 means “no diversity”, and 1 means “infinity diversity”. The Simpson’s Reciprocal Index is slightly different as it considers the number of species besides the abundance of each one, to create a proportion of each species. The Simpson’s Reciprocal Index is mainly used to evaluate a site’s evenness, using the total number of species as a measurement.

In addition, the Bray-Curtis matrix was used to compare the dissimilarity (or similarity) between the species richness and abundance of the two sites. This index is used to quantify the degree of overlap of distribution and composition of species in two or more distinct locations (Féret & Asner, 2014). For the Bray-Curtis matrix, the values go from 0 to 1, 0 signifying that the two sites share all the same species, and 1, meaning that the sites do not share any species. Lastly, a Whittaker plot was done for each study site to evaluate species abundance and evenness. That would help identify each study site’s most common or dominant and rare species.

The Paleontological Statistics (PAST) software was used to analyze the data more accurately and efficiently. It works with spreadsheet-type data entry where users can perform various statistical analysis tests, mainly focusing on paleontological and ecological analysis (Hammer *et al*., 2001).

## Results

East and South campus showed similar values for species richness but not for abundance. East campus showed 31 species of arthropods and 102 individuals, primarily caterpillars, spiders, and ants. Conversely, the South campus exhibited only 29 species but 194 individuals, mainly comprised of ants, ticks, and spiders.

Regarding the diversity indices, Simpson’s Index of Diversity for the East campus was 0.9433 and 0.8977 for the South site. Also, the D values for the Simpson’s Reciprocal Index for the East and South sites were 17.63389831 and 9.770509, respectively. However, the evenness value for the East campus was 0.568835429, while the South campus’s was 0.336914.

The Shannon’s indices followed a similar trend: the East campus displays a more excellent value of 3.119, while the South campus has a value of 2.688. In addition, the Bray-Curtis Index was determined for both sites, resulting in a value of 0.76923077, or a 76.923077% similarity between sites.

The lower and upper confidence levels for the Simpson’s Index of Diversity were 0.9275 and 0.9529 for the East site. Contrarily, the South site’s lower and upper confidence levels were 0.8733 and 0.9177, respectively. The Shannon’s Index’s lower and upper confidence levels were calculated. The East site had a lower confidence value of 3.006, while the upper confidence value was 3.225. On the South side, the lower confidence value of the Shannon’s Index was 2.594, while the upper confidence value was 2.839.

Fig 1 expresses the Whittaker plot that contains the species richness and relative abundance of the East site, whereas Fig 2 proves the Whittaker plot of the richness and relative abundance of the South site.

**Fig 1.**
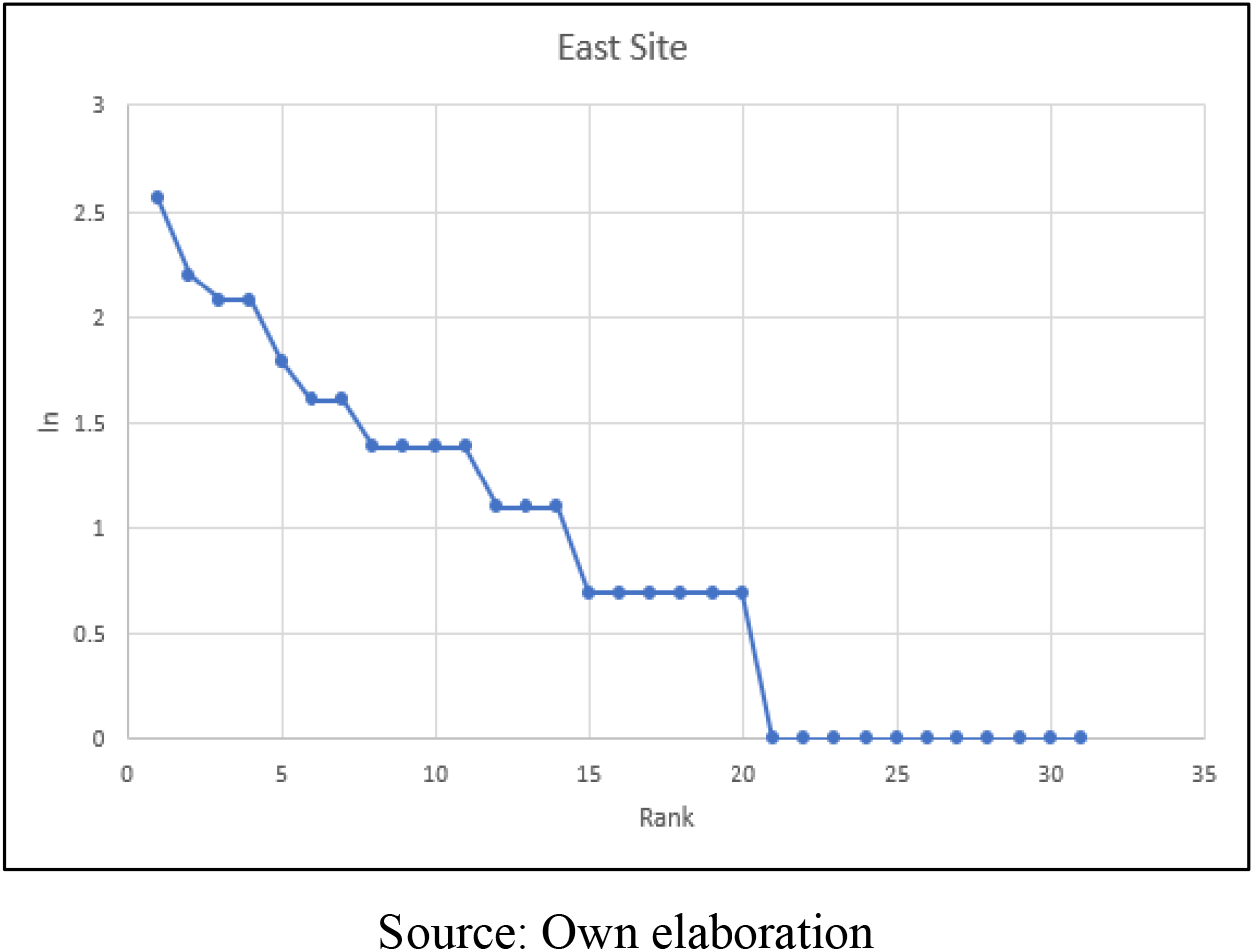
Whittaker plot of the species richness and relative abundance of the East site

**Fig 2.**
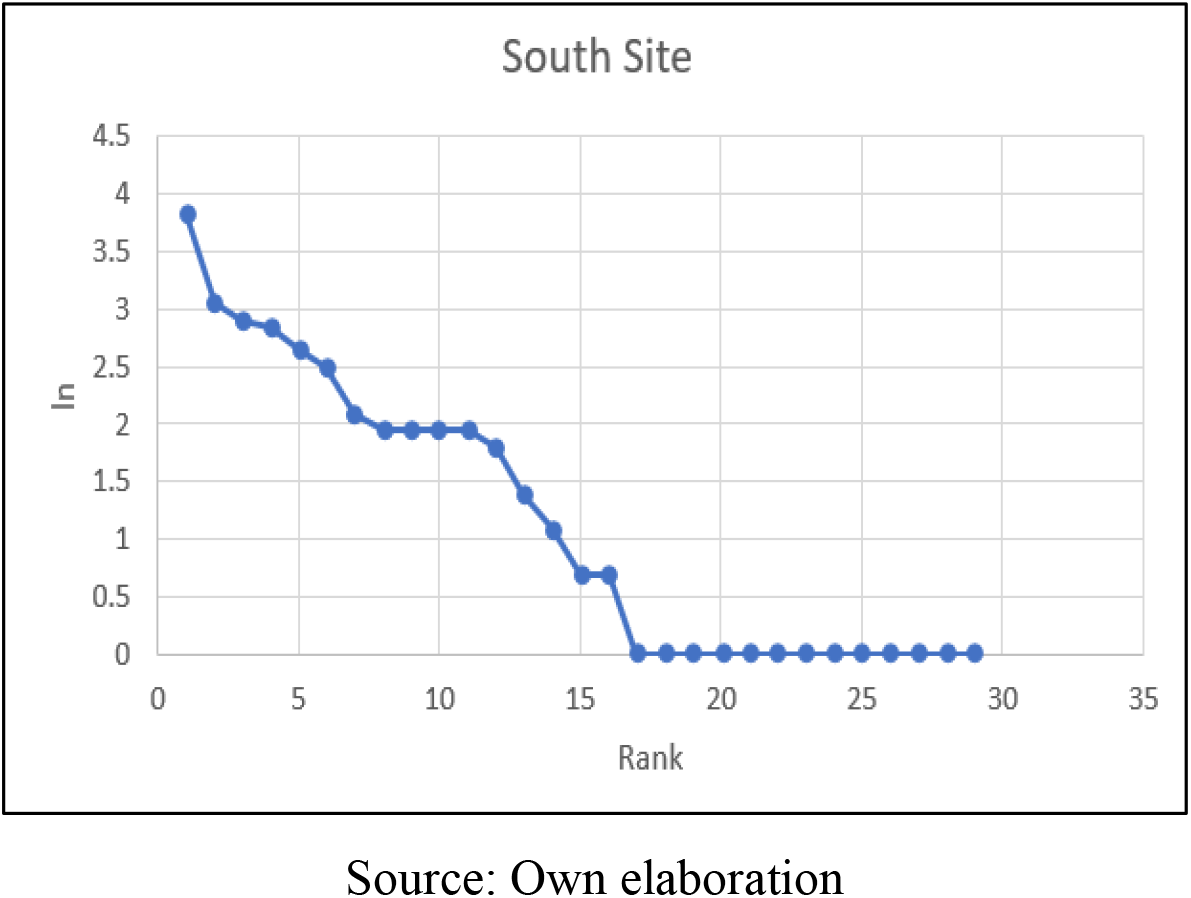
Whittaker plot of the species richness and relative abundance of the South site

## Discussion

The East site, the site with habitat destruction, only has 102 recorded arthropods. In contrast, the South site showed a notable increase in the abundance of arthropods, with 194. The conclusion is clear: humankind has contributed to the arthropods’ risk of extinction. Nonetheless, the Shannon’s Index value for the East side was slightly higher than that of the South side, with a difference of around 0.431. The evenness values were also calculated from the Shannon’s Index for both study sites.

For these, the natural logarithm of the total number of species recorded in that site was used: 31 for the East campus and 29 for the South campus. The resulting values were 0.908274 and 0.798267, respectively. According to these values, the arthropods in the East area are highly evenly distributed, and while the arthropods in the South area are considerably evenly distributed, the ones in the East are about 10% more evenly distributed. According to Shannon’s Diversity Index, the arthropod community located East of the TAMIU campus has a more biologically diverse ecosystem.

Like the Shannon’s Index, the Simpson’s Diversity Index showed that the East site had a value of 0.9433, and the South site had a value of 0.8977. Both locations displayed a high diversity according to the D-values. However, the site located East of the TAMIU campus shows a superior value, close to a 4.5% difference. In the case of the East campus, there is a 94% chance that two randomly selected individuals will belong to different species, meaning there is a high degree of evenness in the area. This value is reasonably close to 100% (or 1.0), and considering the lower and upper confidence values, it can only strengthen the high evenness of this area. Similarly, the South campus displays high evenness but is considerably lower in comparison.

Simpson’s Reciprocal Index values showed a similar trend to those of the previous indices. The East site, having a value of 17.62, is the highest of the two sites. Also, using the total number of species recorded on that site, the evenness value was calculated, resulting in a value of 0.568835429. For the South side, the resulting values were 9.770509 and 0.336914, respectively. This index considers both species richness, abundance, and evenness in the community, suggesting low diversity in both study sites. Unlike the previous indices, Simpson’s Reciprocal Index indicates a low diversity of arthropods in these sites, which had high or at least adequate diversity and evenness according to the other indices.

Additionally, the Bray-Curtis Index indicates a value of 0.76923077, comparing the East and South sites. This value implies a moderately high similarity between the species richness and abundance of the two communities. It takes into account the 14 species found in both habitats, as well as the difference in abundance of individuals. However, considering the two sites are relatively close (∼3 km), they are dissimilar. Although most arthropods recorded were non-flying, different species of spiders and ticks can travel even longer distances through different means (Bram & George, 2000).

Lastly, the Whittaker plots (Figs. 1 and 2) show a relatively uneven abundance distribution for both the East and South locations. In the case of the East site, the species are spread from 2.56 to 0 (minimum abundance of 1). A portion of the species is evenly distributed at the bottom of the graph, with only one individual recorded. These eleven species are evenly distributed; however, they only represent about 35% of the arthropod community in the East site.

In addition, these 11 species are rarely found in this area, as they were only recorded once throughout the study. In contrast, other dominant species were found several times throughout the study, such as the longhorn crazy ant (*Paratrechina longicornis)* or fall armyworm (*Spodoptera frugiperda*) caterpillars, which were recorded 13 times in the East site.

These two species alone correspond to over 20% of the site’s total abundance, with 31 species recorded. This demonstrates a low evenness of the site, similar to the one shown by the Bray-Curtis matrix. Likewise, the Whittaker plot of the South site shows a disproportionate species richness. This one shows 13 species at the bottom of the graph, with a value of 0 (abundance of 1), representing close to 45% of the total species recorded (29) in the location.

While on the other end of the graph, there are species with values above 3 (abundance of 21+). The two most common species were the Eastern black-legged tick and the longhorn crazy ant, with abundances of 46 and 21, respectively. These numbers are equivalent to nearly 35% of the total arthropod community recorded on this site. In comparison, the eleven most rare species account for only 6.7% of the total abundance of the site. That demonstrates the low evenness of the site, where the great majority of the abundance is focused on a few noticeably common species.

## Conclusion

The site with the degraded ecosystem (East site) did not show a clear disadvantage regarding the diversity of arthropods. The species richness of both locations was similar; however, the abundance of arthropods differed significantly from one site to the other, with the southern site having a greater abundance. Additionally, the diversity indices supported the higher diversity and evenness of the East community. Even though Simpson’s Reciprocal Index indicated that both sites had low diversity and evenness, the values of the East campus were still superior to those of the South.

One possible explanation for the difference in arthropod species richness and abundance might be the distribution of mammals, such as white-tailed deer and white-footed mice (*Peromyscus leucopus)*, and host species for other animals, such as ticks. The mammals might leave the East campus due to human disturbance and enter the more unoccupied South campus. This bias might increase certain arthropods’ niches and decrease others’ niches. Another factor influencing arthropod diversity is the plant community in the sites. Similarly, white-tailed deer might influence the plant community by grazing more on the South site rather than the East. That means some plant species essential for certain arthropods might not be available, disrupting the community’s equilibrium.

From a research perspective, this study provides more insight into the arthropod community in the South Texas region, which is an empirical call for help with land and ecosystem management. By understanding more about the region’s ecology, more accurate measurements can be taken to maintain a sustainable ecosystem, considering the critical role of arthropods. These are not only important for pollination but also crucial for disease control and, hence, public health.

